# Predicting Brain Age Using Structural Neuroimaging and Deep Learning

**DOI:** 10.1101/497925

**Authors:** Yogatheesan Varatharajah, Sujeeth Baradwaj, Atilla Kiraly, Diego Ardila, Ravishankar Iyer, Shravya Shetty, Kai Kohlhoff, for the Alzheimer’s Disease Neuroimaging Initiative

**Affiliations:** Department of Electrical and Computer Engineering, University of Illinois at Urbana-Champaign, Urbana, Illinois 61801. Email: {, }; Google LLC., MountainView, California 94043.Email: {,,,, }

## Abstract

Early detection of age-related diseases will greatly benefit from a model of the underlying biological aging process. In this paper, we develop a brain-age predictor by using structural magnetic resonance imaging (SMRI) and deep learning and evaluate the predicted brain age as a marker of brain-aging in Alzheimer’s disease. Our approach does not require any domain knowledge in that it trains end-to-end on the SMRI image itself, and has been validated on real SMRI data collected from elderly subjects. We developed two different models by using convolutional neural network (CNN) based regression and bucket classification to predict brain ages from SMRI images. Our models achieved root mean squared errors (RMSE) of 5.54 and 6.44 years in predicting brain ages of healthy subjects. Further analysis showed that there is a substantial difference between the predicted brain ages of cognitively impaired and healthy subjects with similar chronological ages.

## 1 Introduction

Age-related disease and disability impose a growing burden on society. Since aging effects are subject-specific, markers of the underlying biological aging process are needed to identify people at increased risk of age-related physical and cognitive impairments. Structural MRI images are extremely useful in measuring age-related changes in the brain [1]. Thus, the goals of this study are to develop a brain-age-predicting algorithm based on SMRI images of healthy individuals and to investigate the predicted *brain age*(a person’s age, as predicted from their SMRI image) as a biomarker of age-related diseases using the SMRI images of individuals with cognitive impairment.

The major differences between our approach and previous efforts on this subject are, 1) our use of deep-learning to learn relevant features from SMRI images without requiring domain knowledge, and 2) our validation of the proposed approach with data collected from subjects at risk of developing Alzheimer’s disease, a major aging-related disease. Prior studies [2, 3] have proposed machine-learning-based approaches that use a Gaussian process regression to predict brain age from SMRI images. However, those approaches have relied on features derived from domain knowledge of the structure of the human brain. On the other hand, the authors of [4] have proposed a CNN-based architecture that uses minimal domain information to predict brain age. However, that study was performed using SMRI imaging data collected from children and it is unclear whether it can predict aging-related disease-risk of elderly patients.

We employed a transfer-learning approach using a pretrained Inception-V1 [5] based 3D feature extractor and built a CNN-based model to predict brain age from SMRI images. This model did not require any domain knowledge and predicted brain ages in two different ways, 1) using a regressor, and 2) using a bucketed classifier (described in Section 2). We evaluated our approach using the Alzheimer’s Disease Neuroimaging Initiative (ADNI) dataset [6]. Regression and bucketed classifier methods achieved RMSEs of 5.54 and 6.44 (years) in predicting brain age of cognitively normal subjects. In addition, further analysis showed that the predicted brain ages of cognitively impaired subjects are on average 3.24±2.12 and 2.67±2.35 years higher than their chronological ages when regression and bucketed-classification approaches, respectively, are utilized. In essence, our approach utilizes a CNN-based model for predicting brain age based on SMRI images without using any domain knowledge and demonstrates that brain ages predicted by using our approach can be used to identify age-related disease risk.

## 2 Methodology

The overall flow of our approach is shown in Figure 1. 3D SMRI images of the same physical dimensions were fed as inputs to the pipeline. A 3D feature extractor was utilized to extract feature maps from the input images. Then, a fully connected layer was utilized to make age predictions as a regression task or a bucketed-classification task.

**Figure 1:**
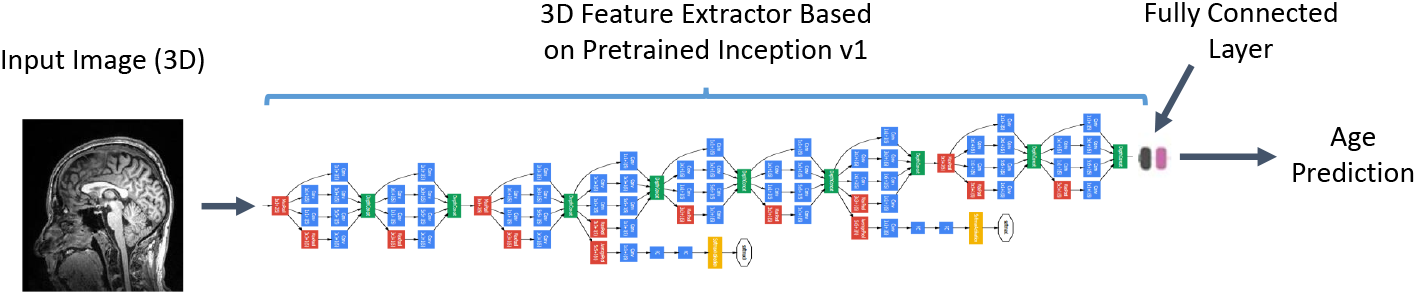
Deep-learning model for predicting brain age (image adapted from [5]).

### Data and preprocessing

We utilized the data collected in the Alzheimer’s Disease Neuroimaging Initiative (ADNI) study [7] to validate our approach. ADNI is an ongoing longitudinal study that periodically collects imaging and blood biomarkers from elderly subjects who are at risk of developing Alzheimer’s disease (AD) with the goal of studying disease progression. We analyzed 12,988 SMRI images of 1484 unique participants for whom ground truth information on their clinical stages of AD and chronological ages was available from the dataset. Clinical stages of AD consist of cognitively normal (CN), mild cognitive impairment (MCI), and Alzheimer’s dementia (AD), and the ages are real numbers between 50 and 100. At the time of their first SMRI scan, 450, 788, and 246 subjects were in the CN, MCI, and AD stages respectively. Multiple SMRI images and corresponding ground truth information were acquired from some of the subjects at different time points. The 3D SMRI images in this dataset were obtained using the MPRAGE sequence [8]. Before performing model training, we resized the raw SMRI images to the same physical dimensions (voxel size), equalized their histograms, and cropped them to a shape of 256 × 256 × 256 (voxels) around the center.

### Model description

A I3D (inflated Inception-v1) feature extractor as described in [9] was utilized to extract feature maps from the input images. One fully connected layer was used to generate age predictions. We predicted brain ages using a regressor (with real number predictions) and a bucketed classifier. In the bucketed-classification approach, the true ages of the subjects were binned into discrete ranges, and the neural network was used to predict the bin to which an SMRI image belonged. The center of the predicted bin was considered as the age predicted by this approach. In our approach, we binned the ages into five buckets, 50–60, 60–70, 70–80, 80–90, and 90–100, and assigned them class labels in {1, 2, 3, 4, 5}. These class labels were used during model training with a cross-entropy loss. The CNN-based models were trained using SMRI images of healthy subjects only, i.e., the training and tuning sets included only cognitively unimpaired subjects. We took this approach to develop a baseline age predictor under healthy aging. Then, the trained models were used to predict the brain ages of a mixture of cognitively impaired and unimpaired subjects. We included some cognitively unimpaired subjects in the testing set to maintain an unbiased testing sample. During model training, we utilized a fivefold cross-validation and patient-based stratification with 50% training, 25% tuning, and 25% testing fractions of the input dataset. Patient-based stratification was utilized to ensure that SMRI images of the same patient never appeared in more than one of the three datasets. In addition, the tuning and testing sets included one SMRI image per subject (their first scan) while the training set included all images available from a subject.

### Evaluation

First, we evaluated model fit using the root mean squared error (RMSE) metric achieved on the tuning set averaged over the five-fold cross-validation. Second, we evaluated the differences in brain age predictions between cognitively unimpaired and impaired subjects in the testing set. We performed that analysis using only the subjects in the testing dataset, and subjects at the MCI and AD clinical stages were considered to be cognitively impaired. We grouped the real-numbered predicted ages of the regression model based on the same ranges used for the bucketed-classification to come up with predicted age groups (see Figure 3a). Note that we have excluded subjects with true ages > 90 in figures 2 and 3 due to privacy concerns and excluded subjects with predicted ages < 60 in figure 2 since there were too few data points.

**Figure 2:**
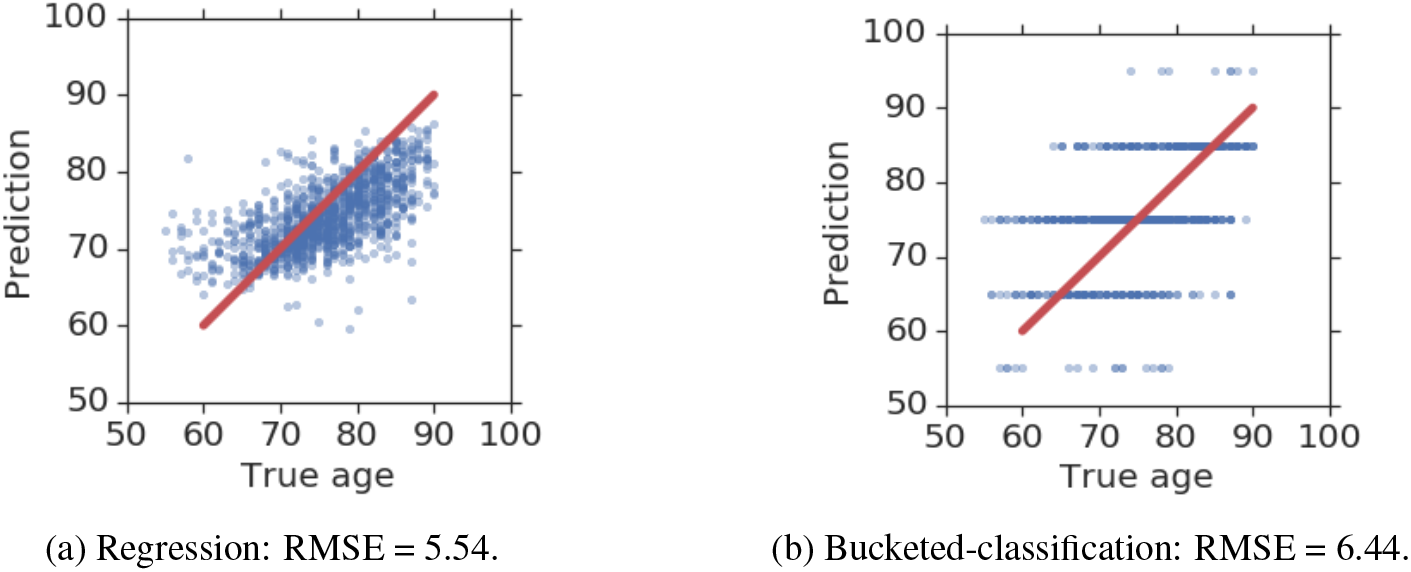
Scatter plots of the predicted ages and true ages of cognitively unimpaired subjects. The red lines indicate the 45° slope on which the ideal predictions should have lain.

**Figure 3:**
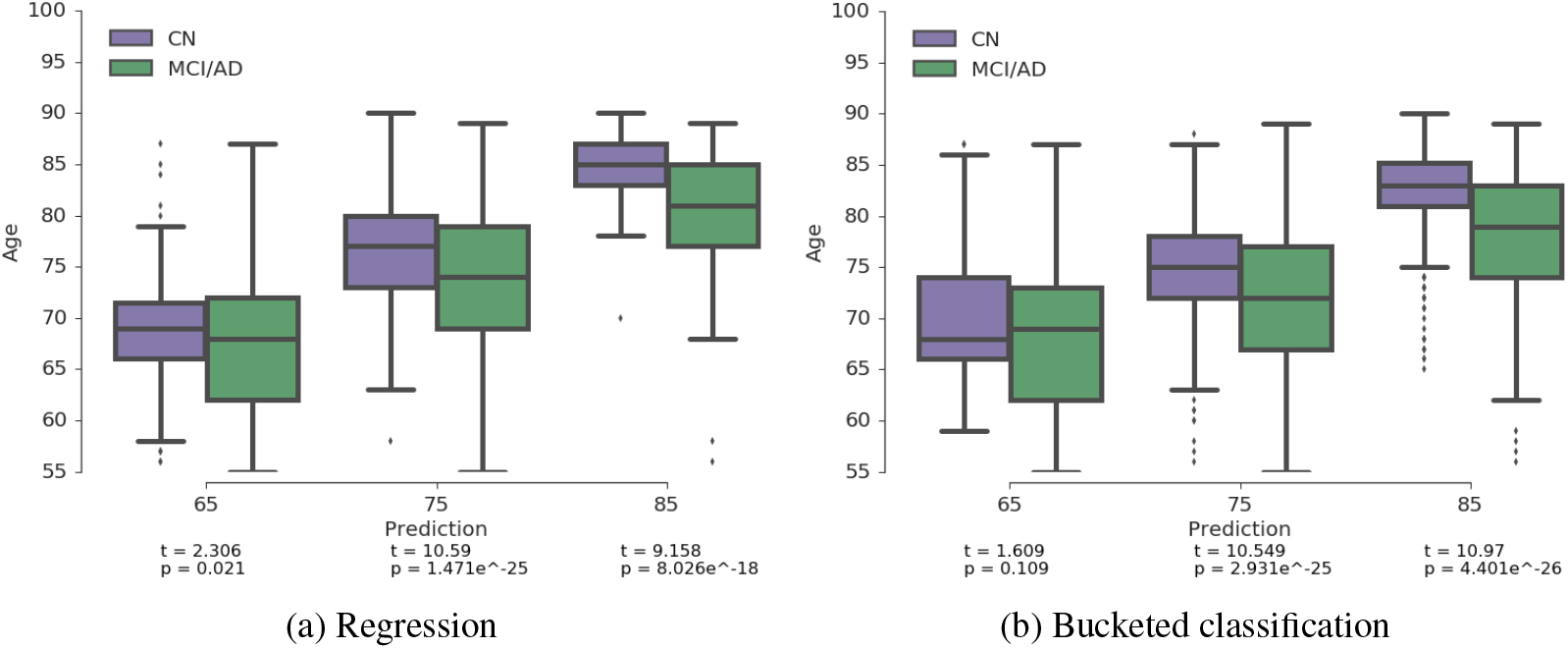
Box plots showing the distributions of true ages (y-axis) for cognitively unimpaired and impaired subjects in the same predicted age group (x-axis). MCI and AD are two disease states of cognitively impairment, while CN is a cognitively normal state.

## 3 Results

Figures 2a and 2b show scatter plots of predicted ages against true ages of cognitively normal subjects in the tuning dataset for the regression model and bucketed-classification model, respectively. The regression model achieved an RMSE of 5.54 while the bucketed-classification model achieved an RMSE of 6.44. Figures 3a and 3b show box-plots of the distributions of true ages for cognitively unimpaired and impaired subjects in the same predicted age group. The figures show that the distribution of the true ages of cognitively impaired subjects consistently has a lower mean value than that of cognitively unimpaired subjects when they were predicted to be in the same age group. Statistical comparisons using independent t-tests indicate that the distributions are sufficiently different in all but one age group (see figure for t-statistics and corresponding p-values). In addition, the predicted ages of cognitively impaired subjects were on average 3.24±2.12 and 2.67±2.35 years higher than their chronological ages in the regression and bucketed-classification approaches respectively.

## 4 Conclusion

We developed two different convolutional neural network (CNN) based approaches based on the transfer-learning paradigm to predict brain ages from SMRI images. Our two models achieved RMSEs of 5.54 and 6.44 (years) in predicting brain ages of cognitively unimpaired subjects. Further analysis showed that there is a substantial difference between the predicted brain ages of cognitively impaired subjects and normal subjects who belong to the same chronological age group. Although our results show that predicted brain ages can identify individuals with age-related diseases, whether they can be used to identify young individuals who are at higher risk of developing age-related diseases later in their lives remains to be seen. In future work, we will obtain additional data to evaluate this hypothesis and perform model optimization to improve prediction performance.

## Acknowledgements

Data used in preparation of this article were obtained from the Alzheimer’s Disease Neuroimaging Initiative (ADNI) database (adni.loni.usc.edu). As such, the investigators within the ADNI contributed to the design and implementation of ADNI and/or provided data but did not participate in analysis or writing of this report. A complete listing of ADNI investigators can be found at: http://adni.loni.usc.edu/wp-content/uploads/how_to_apply/ADNI_Acknowledgement_List.pdf

